# *Pde10a* gates light responses in the SCN to regulate circadian photoentrainment

**DOI:** 10.1101/2025.07.14.664763

**Authors:** R Komal, MB Thomsen, Q Tang, S Yang, H Wang, H Zhao, S Hattar

## Abstract

Light is the principal cue for synchronizing the circadian clock. A common feature of the clock among all organisms is the lack of responsiveness to light during the daytime. To understand the interaction between the circadian clock and light, we described the transcriptome of the suprachiasmatic nucleus (SCN) in mice across different circadian times under both constant darkness (DD) and in response to light exposure. In addition to classifying 10 distinct molecularly-defined SCN neuronal subtypes, we uncovered that SCN exhibits significant transcriptomic responsiveness to light during daytime, the so-called behavioral “dead zone”. We further identified *Pde10a*, a cyclic nucleotide phosphodiesterase, as the first critical component for gating SCN responsiveness to light across the day and thus maintaining robust daily oscillations under regular light-dark conditions.

## INTRODUCTION

Life on earth has evolved to synchronize organismal activity with cyclic changes in the environment, which are driven primarily by the solar day. In mammals, this adaptation is reflected in the daily rhythmic fluctuations in metabolism, physiology, and behavior, which are governed by the central circadian pacemaker located within the suprachiasmatic nucleus (SCN) of the hypothalamus(*1–4*). The SCN maintains an intrinsic circadian clock of approximately 24 hours, which is achieved via a complex network of molecular (transcriptional, translational, and post-translational) oscillators(*3, 4*). To align its intrinsic molecular clock with the external environment, the SCN receives direct photic input from the retina(*5*), which enables it to synchronize with the 24-hour solar day through a process known as photoentrainment(*6, 7*). The molecular machinery of the clock is tightly regulated by intracellular signaling pathways(*4, 8*). Recent studies highlight cAMP as a central regulator within the SCN, crucial for initiating, sustaining, and reinforcing its rhythmic activity(*9, 10*).

The influence of light on the SCN clock, however, is not uniform, and varies considerably depending on the circadian phase during which the light exposure occurs(*11, 12*). The acute effects of light on the circadian clock have been studied by maintaining organisms in constant darkness (DD), where they rely solely on their intrinsic clock to regulate behavior in the absence of external cues, and exposing them to light pulses. Under these conditions, light exposure significantly affects the SCN clock during the subjective night (the animal’s internal circadian nighttime), but not during the subjective day (the animal’s internal circadian daytime)(*11*). In the early night, a light stimulus induces a phase delay of the clock, while in the late night, the same light causes a phase advance(*11*). During the subjective day, however, light exposure does not elicit any changes in the clock phase, regardless of intensity or duration. This time window when the SCN clock is insensitive to the light input is referred to as the “dead zone”(*6, 12–14*). The daily cycle of SCN light sensitivity provides a critical mechanism for organisms to synchronize their internal rhythms with external environmental cues; selective responsiveness of the clock ensures that light can reset circadian timing during the night, particularly when environmental changes such as dawn and dusk serve as reliable signals(*7*). During the day, the clock remains insulated from natural light fluctuations, preventing maladaptive phase shifts that could arise from moving between different light environments, such as from shade or burrows to direct sunlight. This daytime immutability is vital for maintaining stable circadian rhythms, as it safeguards physiological and behavioral processes essential for fitness and survival by preventing constant disruptions that would otherwise compromise the clock’s alignment with the solar cycle.

Recent transcriptomic studies have shed light on the molecular mechanisms underlying the SCN’s response to light cues, particularly during the early night(*15, 16*). Despite progress in elucidating signaling pathways for light-induced responses(*14, 17–19*), the mechanisms by which the circadian clock responds differentially to light during different times of the day, and particularly mechanisms that prevent the SCN clock from responding to light during the “dead zone”, remain poorly understood.

Here, we employed deep single nucleus transcriptome profiling of the mouse SCN to comprehensively characterize the SCN’s light response across different circadian times – during subjective day, early subjective night, and late subjective night. This approach allowed us to classify 10 distinct molecularly defined SCN neuronal subtypes. Notably, contrary to previous assumptions of transcriptional unresponsiveness, we observed a robust light-induced response during the subjective day, the so-called behavioral “dead zone”. This analysis allowed us to identify *Pde10a,* a cyclic nucleotide phosphodiesterase that regulates cAMP second messenger levels, as the first critical component for gating the SCN responsiveness to light across the day and an essential factor for maintaining robust circadian oscillations under regular light-dark conditions.

## RESULTS

### Circadian and light-dependent transcriptional annotation and spatial distribution of SCN neuronal subtypes

Circadian behavioral responses to light exposure vary by time of day: a refractory period, referred to as the ‘dead zone,’ occurs during daytime hours, while phase delays and phase advances are induced during the early and late night, respectively(*6, 11, 12*). To understand the molecular mechanisms by which the circadian clock responds to light during different times of the day, we conducted single nucleus RNA-sequencing of the mouse anterior hypothalamus following light exposure at three key time points: Circadian Time 6 (CT6, mid-day), CT14 (early night), and CT22 (late night). At each time point, animals housed in constant darkness (DD) were either exposed to 15 minutes of bright white light (L) or kept in darkness (D), and hypothalamus tissue containing the SCN was collected 45 minutes after exposure to allow for peak transcriptional activation(*16*) (Fig. 1A). From all six time points, we isolated a total of 148,806 nuclei (27,348 CT6D; 28,510 CT6L; 23,185 CT14D; 17,003 CT14L; 34,290 CT22D; 18,470 CT22L) with an average of 1,845 genes detected per nucleus, (Fig. S1A, Data S1).

**Fig. 1.**
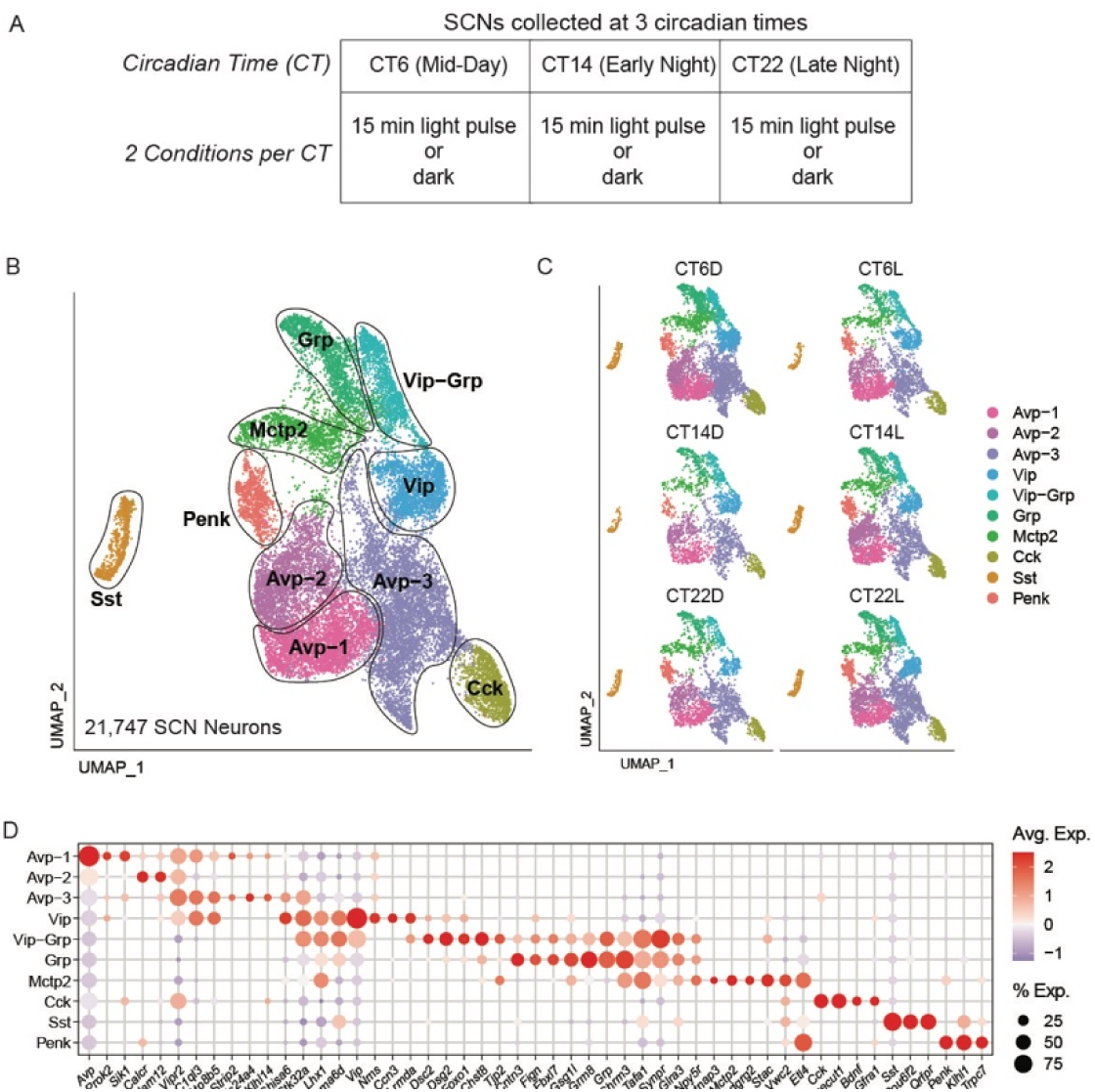
Transcriptomic characterization of SCN neurons under constant darkness and in response to light exposure across three circadian phases. See also Fig. S1. **(A).** Experimental design. Mice housed under 12L:12D cycles were kept in darkness for one day. After that, hypothalamus tissues containing suprachiasmatic nuclei (SCN) were collected at each of three circadian times (CT): CT6 (mid-day), CT14 (early night), or CT22 (late night). For each circadian time, mice were either kept in dark or exposed to a 15-minute light pulse. SCNs were micro-dissected after 45 minutes of the light exposure. Nuclei were then extracted and subjected to single-nucleus RNA sequencing. Note that each time point corresponds to a different behavioral response to light exposure: no phase shift at CT6, phase delay at CT14, and phase advance at CT22. **(B).** UMAP visualization of unsupervised clustering of 21,747 SCN neurons reveals 10 distinct neuronal subtypes based on gene expression profiles. Each cluster is represented by Avp-1, Avp-2, Avp-3, Vip, Vip-Grp, Grp, Mctp2, Penk, Sst, and Cck, respectively. **(C).** UMAP plots of SCN neuronal subtypes across different circadian times and light conditions: CT6D (dark), CT6L (light), CT14D (dark), CT14L (light), CT22D (dark), and CT22L (light). Each plot contains the same 10 neuronal subtypes as those in panel B, suggesting the conservation of neuronal subtypes across circadian times and light conditions. **(D).** Dot plot showing marker gene expression patterns (labeled on the x-axis) of 10 SCN neuronal subtypes.

SCN neurons were identified among subpopulations of hypothalamic GABAergic neurons based on the expression of known SCN-enriched transcripts(*15, 20, 21*) such as *Avp*, *Vip*, *Nms*, and *Lhx1* (see Materials Methods; Fig. S1B). We used anchor-based integration and unsupervised clustering to identify 10 distinct subtypes of SCN neurons (Fig. 1B) which were present across all circadian phases and light conditions (Fig. 1C, Data S1). Marker gene enrichment in SCN neuronal subtypes revealed that many could be delineated by neuropeptide expression (Data S2). 9 out of 10 subtypes were classified based on enrichment of known SCN neuropeptide transmitters(*22, 23*), including *Avp* (Clusters ‘Avp-1’, ‘Avp-2’, ‘Avp-3’), *Vip* and *Grp* (Clusters ‘Vip’, ‘Vip-Grp’, ‘Grp’), *Cck* (Cluster ‘Cck’), *Sst* (Cluster ‘Sst’), and *Penk* (Cluster ‘Penk’) (Fig. 1D and S1F). One subtype was not specifically defined by neuropeptide expression, but was enriched for the transcription factor *Lhx1* and other SCN-specific marker genes (Fig. 1D). We designated this cluster as the ‘Mctp2’ subtype, owing to its exclusive expression of the conserved, calcium-sensitive, synaptic plasticity-related gene *Mctp2*(*24, 25*).

The three Avp-expressing clusters were distinguished by the presence of *Avp*, *Vipr2,* and *C1ql3*. *Avp* and *Vipr2* are well-established in the field, and *C1ql3* is noteworthy as a recognized synaptic organizing protein in the SCN(*26*). The ‘Avp-1’ cluster exhibited the highest *Avp* expression, while the ‘Avp-2’ cluster was discriminated from ‘Avp-1’ by the co-expression of *Calcr* and low expression of *C1ql3*. Interestingly, the ‘Avp-3’ cluster, while the most abundant, exhibited the lowest *Avp* expression, yet maintained expression of other Avp-associated markers such as *Vipr2* and *C1ql3* (Fig. 1D). Despite these distinct features, all three clusters demonstrated a high degree of transcriptional similarity overall, with shared variable gene expression ranging from 86% to 95% (Fig. S1C).

Two distinct clusters, ‘Vip’ and ‘Vip-Grp’, where characterized by high levels of *Vip* expression. Notably, the ‘Vip’ cluster exhibited substantial overlap in gene expression with the three ‘Avp’ clusters (72-88%, Fig. S1C), including enrichment for Avp-associated markers such as *Vipr2*, *C1ql3*, and *Nms* (Fig. 1D). In contrast, the ‘Vip-Grp’ cluster was transcriptionally distinct from Avp cells, with less than 50% similarity (Fig. S1C). This ‘Vip-Grp’ cluster also shared several markers with ‘Grp’ and ‘Mctp2’ clusters, including synapse- and neurotransmission-related genes like *Synpr*, *Glra3*, and *Chrm3* (Fig. 1D). The ‘Grp’ cluster was strongly enriched for *Grp* expression in addition to cell adhesion molecule *Cntn3* and metabotropic glutamate receptor *Grm8* (Fig. 1D). The ‘Mctp2’ cluster was similarly distinguished by the expression of transmembrane, synaptic, and cell adhesion genes, including *Mctp2*, *Cntnap3*, and *Stac* (Fig. 1D and S1F), and exhibited transcriptional similarity to *Vip*- and *Grp*-enriched clusters (87-90%; Fig. S1C).

The ‘Cck’ cluster had high similarity to the ‘Avp’ clusters (79-87%) and was uniquely labeled by *Bdnf* expression as reported previously(*15*) (Fig. 1D and S1C;). While the ‘Penk’ and ‘Sst’ clusters were identified as distinct subtypes, they exhibit 70-84% (Fig. S1C) similarity with the ‘Grp’ and ‘Mctp2’ clusters.

Following the identification of 10 transcriptionally distinct SCN neuronal subtypes, we sought to map their neuroanatomical distribution and determine how these diverse neuronal subtypes align with, or diverge from, the classical ‘core-shell’ architecture of the SCN(*27–29*). To achieve simultaneous visualization of SCN neuronal subtypes, we used multiplexed single molecule fluorescence *in situ* hybridization (FISH) in brain slices collected at timepoints that matched those used for sequencing (CT6, CT14 and CT22, dark and light). We selected marker genes that are enriched, specific, and non-overlapping to visualize each neuronal subtype (see Table 1, Fig. 1D) and found that our selected FISH probes each labeled a unique and spatially defined population of cells in the SCN (Fig. 2).

**Fig. 2.**
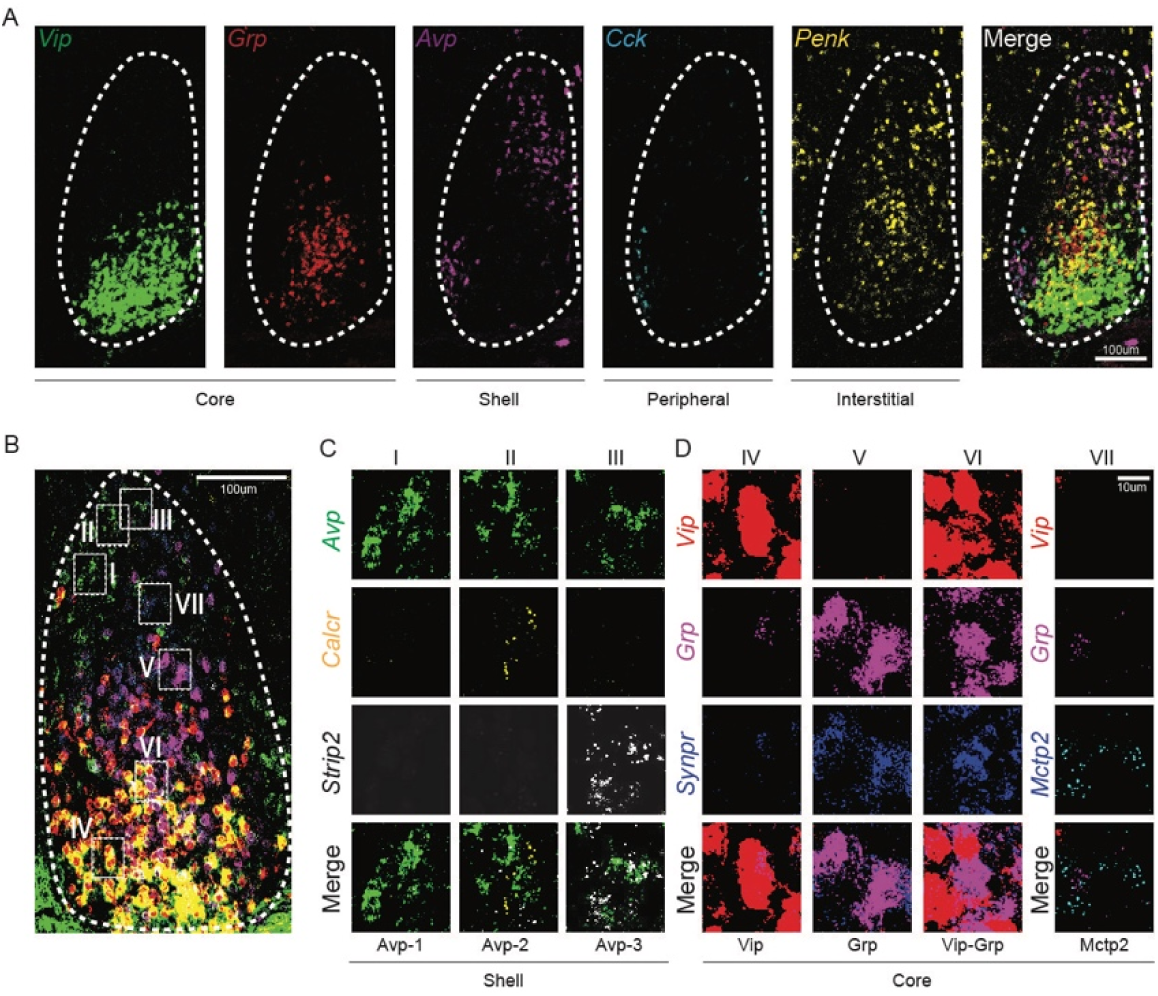
Spatial organization of neuronal subtypes in the SCN. See also Fig. S2. **(A).** 4 different SCN regions (core, shell, peripheral, and interstitial) depicted by the spatial distribution of 5 marker genes with HiPlex (FISH) in the same SCN. From left to right: core region (*Vip;* green, *Grp;* red), shell region (*Avp*; purple), peripheral region (*Cck*; cyan), interstitial region (*Penk;* yellow), and merged. Scale bar: 100 μm. **(B).** Simultaneous visualization of 7 distinct neuronal subtypes (I, II, III, IV, V, VI and VII) in the same SCN by marker gene expression with HiPlex. Scale bar: 100 μm. **(C).** Detailed views of SCN-shell region, showing neuronal subtypes I (*Avp*, green), II (*Avp*, green; *Calcr*, yellow), and III (*Avp*, green; *Strip2*, white). Scale bar: 10 μm. **(D).** Detailed views of SCN-core region, showing neuronal subtypes IV (*Vip*, red), V (*Grp*, purple; *Synpr*, blue), VI (*Vip*, red; *Grp*, purple; *Synpr*, blue) and VII (*Mctp2,* cyan). Scale bar: 10 μm.

**Table 1.**
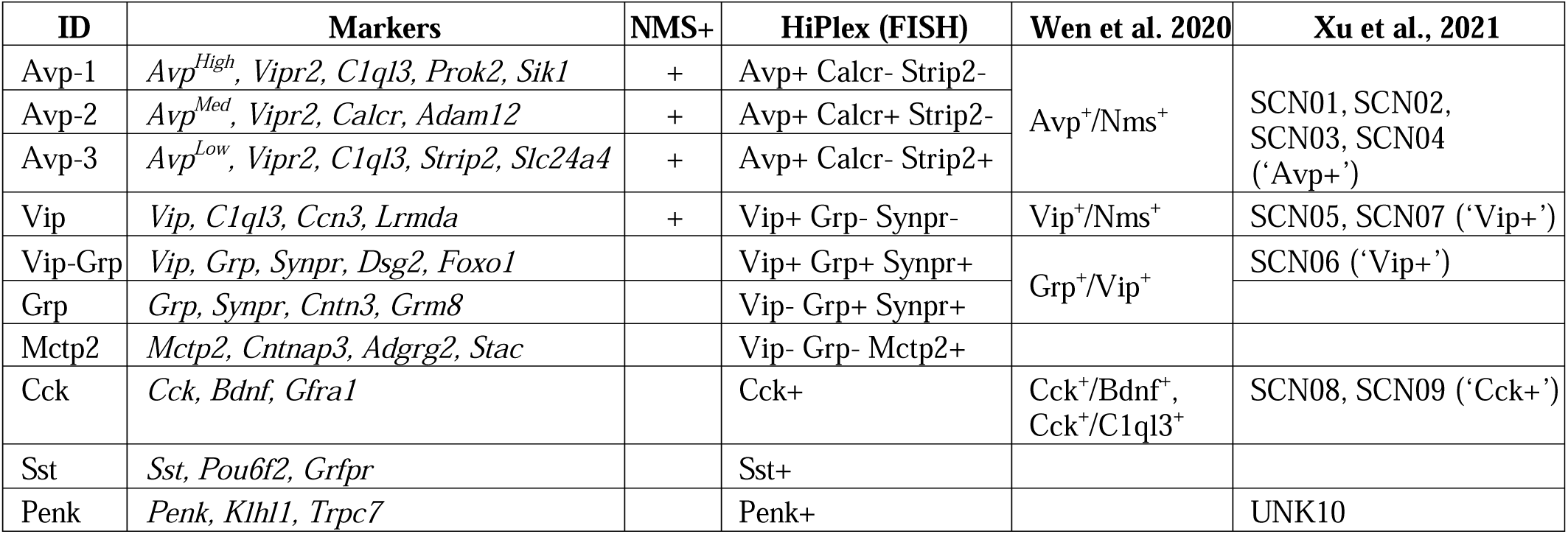
Marker genes to visualize 10 distinct SCN neuronal subtypes, HiPlex assay, and comparison of our SCN neuronal subtypes with previously published dataset. Related to **Figure 1** and **Figure 2.**

Avp+ cells were found in the dorsolateral portion of the SCN, corresponding to the conventional “shell” region (Fig. 2A), and three subsets of ‘Avp’ neurons were distinguished by differential labelling of *Avp*, *Calcr*, and *Strip2* (‘Avp1’: *Avp*+/*Calcr*-/*Strip2*-; ‘Avp2’: *Avp*+/*Calcr*+/*Strip*2-; ‘Avp3’: *Avp*+/*Calcr*-/*Strip2*+; Fig. 2B, 2C and Table 1). *Vip*+ and *Grp+* cells were present in the central/ventromedial portion of the SCN, corresponding to the conventional “core” region (Fig. 2A and 2B). We used *Synpr* to distinguish ‘Vip-Grp’ (Vip+/Grp+/Synpr+) and ‘Grp’ (Vip-/Grp+/Synpr+) neurons from ‘Vip’ (Vip+/Grp-/Synpr-) neurons (Fig. 2B and 2D, Table 1). Mctp2+ cells, although present in the core region of the SCN (Fig. 2B), did not express core genes *Vip* and *Grp* (Fig. 2D).

Cck+, Sst+, and Penk+ cells showed distinct distribution patterns in the SCN. Cck+ cells were found around the anatomical border of the SCN, while Sst+ and Penk+ cells were distributed throughout the SCN (Fig. 2A, S2A, and S2C). Based on these observations, we termed Cck+ cells “peripheral neurons” and Sst+ and Penk+ cells “interstitial neurons” (Fig. 2A and S2B), reflecting their divergence from core/shell organization.

Taken together, we identified 10 transcriptionally distinct SCN neuronal subtypess harmonized across six conditions and organized into four anatomical domains: ‘shell’, ‘core’, ‘peripheral’, and ‘interstitial’. Despite overall transcriptional similarity of SCN neurons, each subtype exhibits distinguishing features *in vivo*, driven by genes related to neurotransmission, cell adhesion, and synaptic function. These molecular differences likely reflect the functional specialization of each subtype and correlate with their distinct spatial distributions within the SCN.

### Dynamic circadian gene expression in SCN neuronal subtypes

Our transcriptional analysis showed that all 10 subtypes of SCN neurons exhibit circadian rhythms in their gene expression (Fig. S3-1A). In general, the magnitude of differential gene expression in each subtype was higher between day and night compared to early vs. late night (Fig. S3-1A). There was a striking dynamic range of rhythmic gene expression, with the number of differentially expressed genes fluctuating from dozens to thousands across subtypes and timepoints (Fig. S3-1A). We confirmed that core clock genes exhibit the expected temporal dynamics and antiphase relationship in dark controls(*30, 31*). In particular, *Per2* peaked at CT6 compared with its late-night levels at CT22 (adjusted p-value < 0.05) in Avp-1, Avp-2, Avp-3, and Vip subtypes (Fig. S3-1C, Data S3), while *Bmal* (*Arntl*) reached its maximum expression at CT14 (adjusted p-value < 0.05 when comparing CT14 vs. CT6, Fig. S3-1D, Data S3). These observations align with the canonical transcriptional-translational feedback loop and indicate that we can reliably detect circadian gene oscillations in specific SCN neuron populations.

We examined the similarity of the transcriptional response between all SCN neuronal subtypes and found that, surprisingly, the sets of differentially expressed genes were largely distinct with a low degree of overlap (59% max overlap) between subtypes at different circadian times (Fig. S3-1B). In general, we noted that ‘Grp’ and ‘Vip-Grp’ neuron responses were more similar than other types, whereas Vip neuron responses correlated more strongly with Avp neurons, further highlighting the distinction between Vip and Vip-Grp subtypes (Fig. S3-1B). We used gene ontology to examine the pathways and components that change throughout the day in SCN neurons and found that many transcripts were involved in synapse structure and neurotransmission regulation, particularly in Avp subtypes (Fig. S1D and S1E), consistent with reports that the cellular clock in Avp neurons of the SCN is crucial for inter-neuronal connectivity and the regulation of circadian behavioral rhythm(*32*).

Next, we used FISH to examine the circadian expression of our selected neuronal subtype marker genes (*Avp, Vip, Grp, Mctp2, C1ql3, Synpr, Mctp2, Strip2,* and *Calcr*) at the same time points used for sequencing. *Avp*, *C1ql3*, *Synpr*, and *Mctp2* showed significant circadian rhythms in their expression levels (Fig. S3-1E). Interestingly, the robust rhythmic expression of *C1ql3* and *Synpr*, known for their involvement in synaptic function(*26, 33*), suggests that the synaptic properties of the SCN change in a neuronal subtype-specific manner throughout the day. In general, these changes were more dynamic than those observed in neuropeptide marker gene expression like *Vip* and *Grp* (Fig. S3-1E). These results are consistent with previous studies demonstrating that daily morphological changes in clock neurons are accompanied by changes in morphology and synaptic connectivity(*34–36*).

### Light induces diverse time-specific transcriptional responses in SCN neuronal subtypes

To better understand the diversity of SCN light responses across different circadian phases, we examined light-induced transcriptional changes in each neuronal subtype (Fig. 3A-3D). Surprisingly, we observed a substantial number of differentially expressed genes in all SCN neuronal subtypes during the behavioral “dead-zone” (CT6) following a light stimulus (Fig. 3A and 3C). Notably, significant upregulation of genes at CT6 (mean: 1571 upregulated DEGs; Fig. 3A) was observed in all SCN neuronal subtypes, whereas substantial downregulation occurs primarily in the “Shell” neuronal subtypes (Avp+ mean downregulated DEGs: 1191; non-Avp mean downregulated DEGs: 17; Fig. 3C). In contrast, the light-induced transcriptional response during the subjective night (CT14, CT22), where phase shifts are observed, was characterized by widespread downregulation of genes, with limited upregulation (mean upregulated/downregulated DEGs: CT14=61/347, CT22=33/236; Fig. 3A and 3C). Previous studies of light’s impact on circadian rhythms have mostly focused on genes that are upregulated in response to light(*4, 37–40*). However, the role of gene repression has often been overlooked. This analysis encourages a broader perspective to better understand the full effects of light on circadian rhythms.

**Fig. 3.**
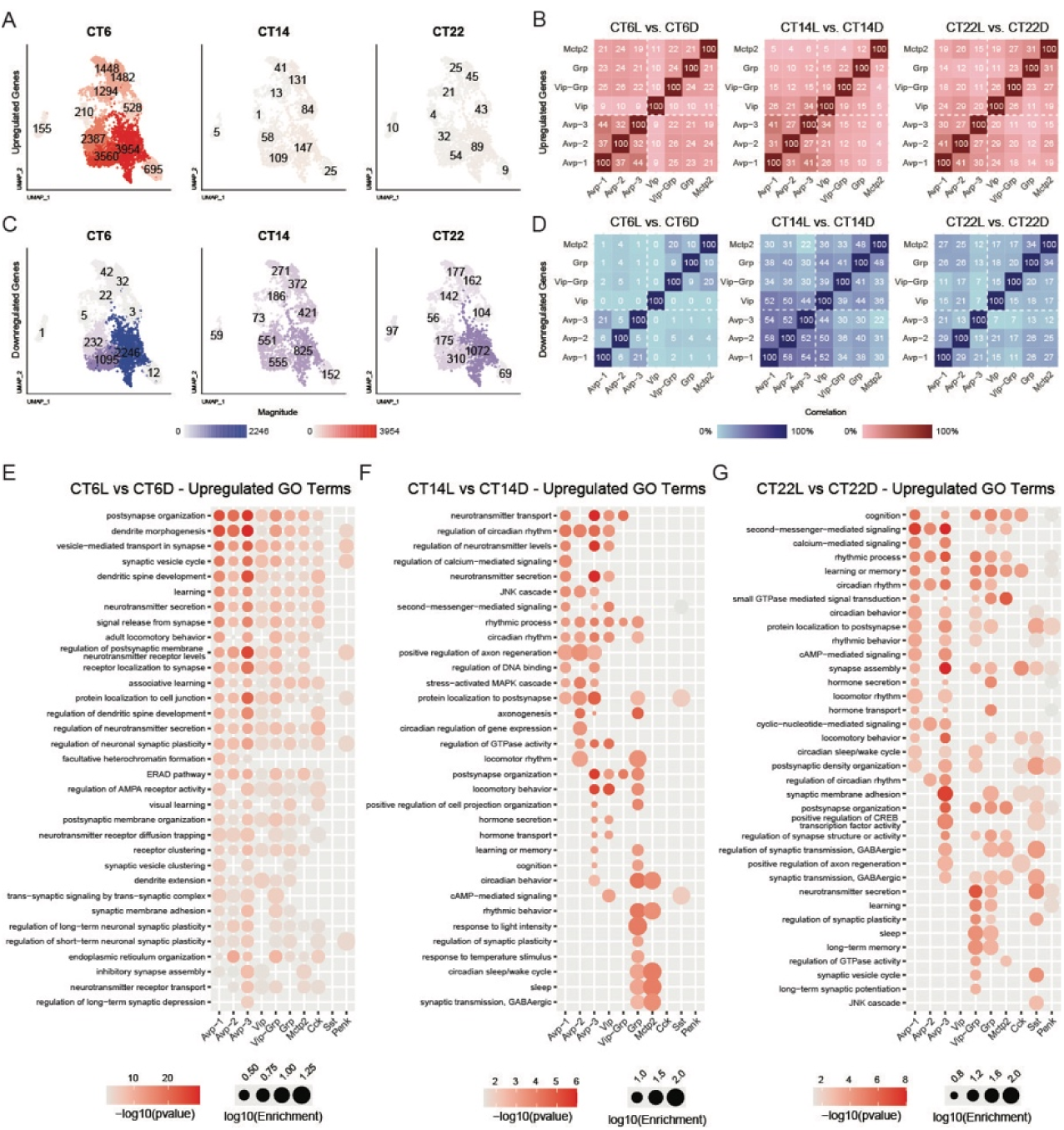
Light-induced transcriptional changes in the SCN at CT6, CT14 and CT 22. See also Fig. S3-1 and S3-2. **(A).** Numbers of upregulated transcripts in each SCN neuronal subtype following a light pulse at three different circadian time: CT6 (left), CT14 (middle) and CT22 (right).10 SCN neuronal subtypes are depicted by UMAP plots. **(B).** Comparison matrices showing the relatedness (Jaccard similarity) of upregulated transcripts among different SCN neuronal subtypes following a light pulse at CT6 (left), CT14 (middle) and CT22 (right). **(C).** Numbers of downregulated transcripts in each SCN neuronal subtype following a light pulse at CT6 (left), CT14 (middle) and CT22 (right). **(D).** Comparison matrices showing the relatedness (Jaccard similarity) of downregulated transcripts among different SCN neuronal subtypes following a light pulse at CT6 (left), CT14 (middle) and CT22 (right). **(E).** Selected Gene Ontology (GO – biological process) term enrichment for light-induced upregulated transcripts (p < 0.05) in each SCN neuronal subtype at CT6. **(F).** Same as in E, but at CT14. **(G).** Same as in E, but at CT22.

As we observed in constant darkness, the transcriptional response to light in each neuron subtype at each time was generally cell type-specific, with low similarity between neuronal subtypes (max 58% overlap) (Fig. 3B and 3D). At CT6, we observed that light upregulated the expression of genes related to synaptic organization and function across nearly all neuronal subtypes (Fig. 3E). This suggests that while light stimuli at this circadian time do not induce phase shifts in the clock, they substantially affect synaptic transmission in the SCN. Consistent with the lack of phase shifts at CT6, fewer upregulated transcripts were related to circadian rhythms. However, at both CT14 and CT22, light upregulated the expression of genes related to circadian behavior, sleep, GABAergic neurotransmission, and synaptic function (Fig. 3F and 3G). In addition, factors related to cAMP second messenger-mediated signaling, calcium signaling, and cAMP Response Element Binding Protein (CREB) transcription factor activity were enriched in response to light uniquely at CT14 and CT22 (Fig. 3F and 3G), consistent with published results(*4, 14, 41*).

A striking downregulation of transcripts related to energy and metabolism was observed at CT14 across all SCN neuronal subtypes (Fig. S3-2A). This reduction was weaker during the late night (CT22), and almost non-existent during the day (CT6) (Fig. S3-2B and S3-2C). Interestingly, the degree of downregulation is correlated with the magnitude of phase responses that the SCN can generate at each respective circadian time, which suggests that metabolic regulation in the SCN might be a key component of light responses.

### *Pde10a* is a rhythmic and light responsive gene in the SCN

Our analyses of dark and light stimuli allow us to identify key genes that are differentially expressed in response to light in SCN neurons at CT6, CT14 and CT22 (Fig. 4A and 4B, Data S3). Consistent with previous studies(*18, 42*), we found that the clock gene *Per2* was upregulated at CT14 in all shell (‘Avp-1’, ‘Avp-2’, ‘Avp-3’) and some core (‘Vip’, ‘Vip-Grp’) subtypes (Fig. 4A and Data S3). Among the differentially expressed genes, Pde10a, which was strongly upregulated in response to light at CT14 and CT22 but not at CT6 in several SCN neuronal subtypes, attracted our attention (Fig. 4A-4C and Data S3. *Pde10a* functions as a cyclic nucleotide phosphodiesterase, which catalyzes the degradation of cyclic nucleotide second messengers(*43, 44*), such as cAMP, cAMP regulates the activity of the transcription factor CREB, which is a key regulator of clock gene expression and phase shifting(*38, 39*). We found that *Pde10a* was broadly and rhythmically expressed in the SCN under constant dark conditions, peaking at CT6, with lower levels at CT22 and minimal expression at CT14. (Fig. 4C). Using FISH in SCN slices, we confirmed that *Pde10a* expression is rhythmic with a peak at CT6 and that its expression increases in response to light only at night (Fig. 4D).

**Fig. 4.**
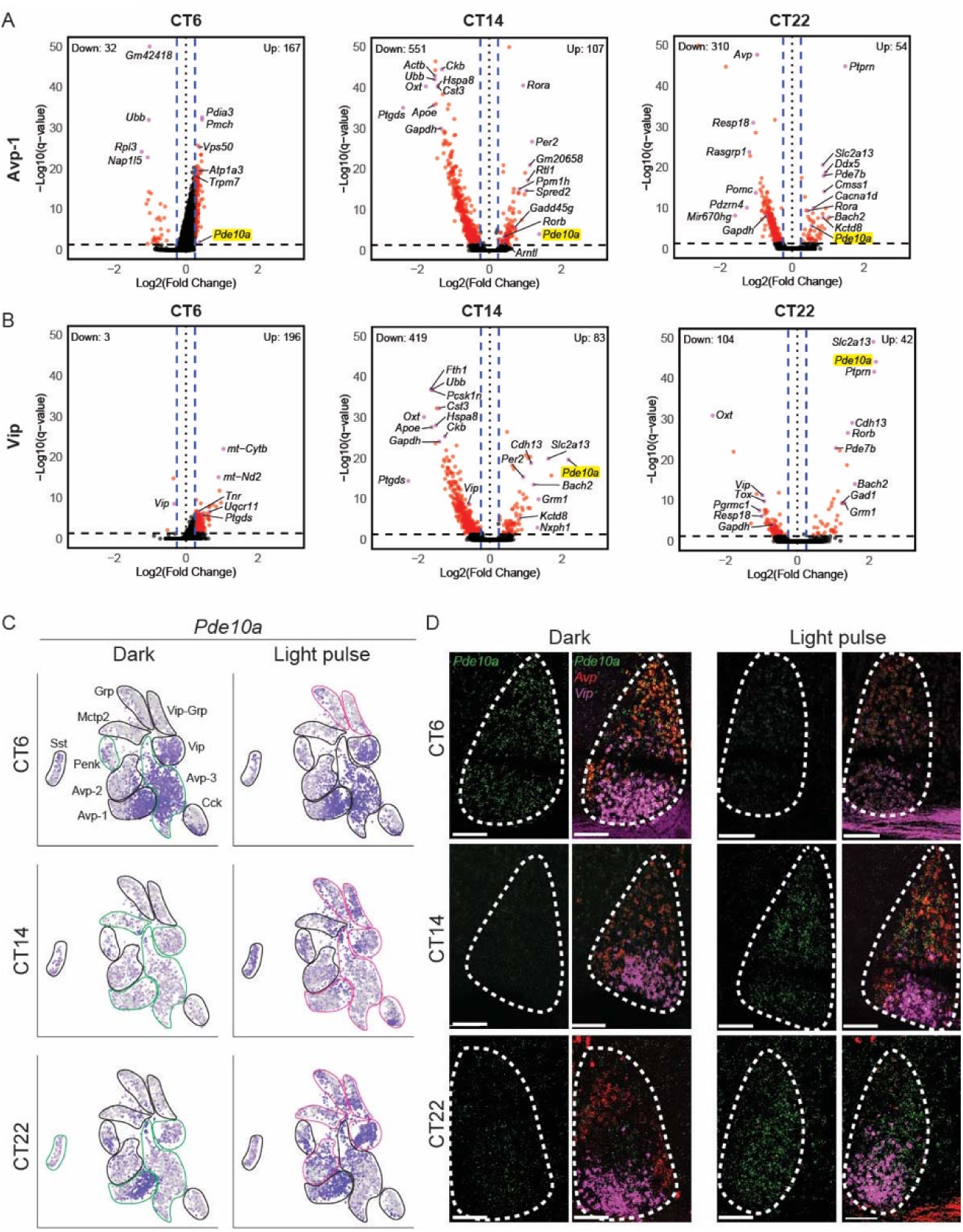
*Pde10a* expression is circadian rhythmic and light responsive in SCN neurons. **(A).** Volcano plots of differentially expressed genes (DEGs) in Avp-1 neuronal subtype following a 15-minute light pulse at CT6 (left), CT14 (middle), and CT22 (right). Red dots: significant DEGs (p < 0.05, |Log2(Fold Change) | > 0.25). *Pde10a* (marked by yellow) is significantly upregulated by light at CT14 and CT22. **(B).** Same as in A, but in the Vip neuronal subtype. *Pde10a* (marked by yellow) is significantly upregulated by light at CT 14 and CT22. **(C).** *Pde10a* expression in SCN neurons depicted by UMAP plots shows that *Pde10a* expression is circadian rhythmic and light responsive. The left panel represents the circadian pattern, and the right panel represents light response at CT6 (upper row), CT14 (middle row) and CT22 (bottom row). *Green circles*: *Pde10a* expression is significantly altered in the circled neuronal subtypes relative to the preceding circadian time (p-adj < 0.05). *Magenta circles*: *Pde10a* expression is significantly upregulated in the circled neuronal subtypes following light pulse (p-adj < 0.05). **(D).** Fluorescence *in-situ* hybridization (FISH) of *Pde10a (*green), *Vip* (magenta) and *Avp* (red) within the SCN. The left panel represents the circadian pattern, and the right panel represents light response at CT6 (upper row), CT14 (middle row) and CT22 (bottom row). Scale bars 100 µm.

### *Pde10a* knock-out mice show abnormal activity pattern under light-dark cycle

Given that *Pde10a* exhibits rhythmic circadian expression and a dynamic light response and that *Pde10a* function is linked to CREB activity(*43, 44*), we used a knock-out mouse model (*Pde10a*-KO) to study the role of *Pde10a* in regulating circadian rhythms and clock responsiveness to light. Animals were placed in individual cages equipped with running wheels and exposed to a regular 12-hour light:12-hour dark (LD) schedule for photoentrainment measurement or in constant darkness for endogenous circadian rhythm measurement. Control mice (*Pde10a^+/+^* and *Pde10a^+/-^*) successfully photoentrained to a LD cycle with a period length of 24 hours (24.0±0.0 hours, n=6 mice; Fig. 5A and S5A), and in DD the average period length was slightly shortened to 23.7 hours as expected (23.7±0.1 hours, n=6 mice; Fig. 5A). In both LD and DD, control mice showed high activity during the early night, progressively decreased activity during the late night, and minimal activity during the daytime (Fig. 5A and 5C). Wavelet analyses(*45*) showed that WT animals have robust amplitude, or strength, of the rhythm in both LD and DD (Fig. 5A). *Pde10a-KO* mice also photoentrained to the LD cycle (23.9±0.1 hours, n=8 mice; Fig. 5B and S5A), but the photoentrainment did not appear as robust as WT animals as illustrated by wavelet analysis (Fig. 5B). Compared to control animals, *Pde10a-KO* mice exhibited an abnormal activity pattern under LD conditions, particularly during transitions from night to daytime (Fig. 5B and 5C). In DD, *Pde10a-KO* mice showed rhythmic wheel running activity with period length (23.7±0.1 hours, n=8 mice Fig. 5B) similar to control mice. Remarkably, *Pde10a-KO* mice showed more robust rhythmic activity in DD compared to LD. These results strongly suggest that *Pde10a* is not required for maintaining the endogenous circadian clock rhythms but is critical for regulating circadian responses to light.

**Fig. 5.**
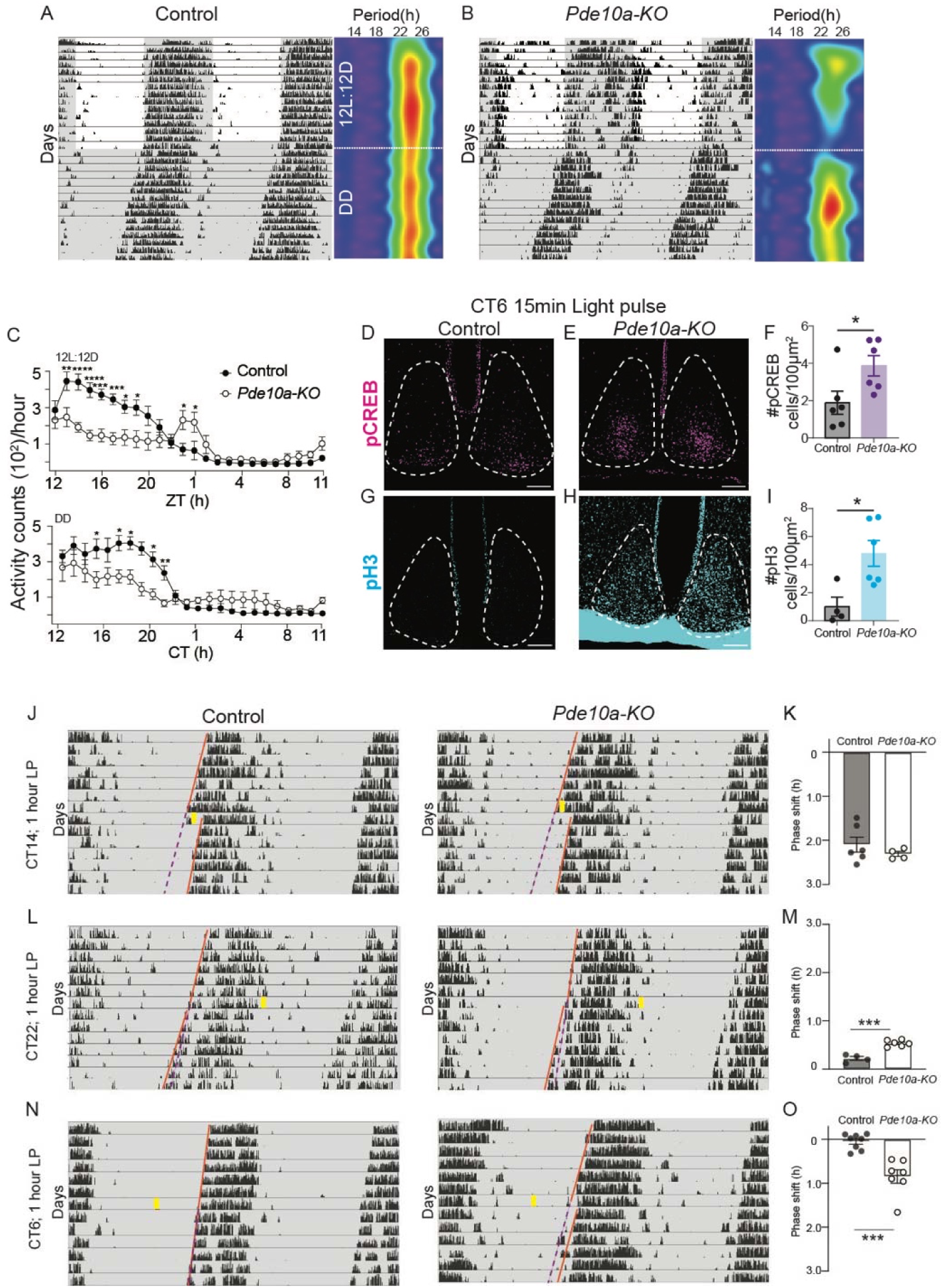
Loss of *Pde10a* results in defective circadian photoentrainment under regular light: dark cycle and permits phase shift of the clock by light in daytime. See also Fig. S5. **A, B and C.** Circadian locomotor activity of control and *Pde10a*-KO mice. **(A).** Left panel, the actogram of a control mouse. Each horizontal line represents one day, with the x-axis showing time and the y-axis indicating successive days. Black bars denote wheel running activity. White background indicates light hours, and grey background signifies dark hours. The mouse was first housed in a regular 12-hour light:12-hour dark (12L:12D) cycle from day 1-14 and then housed in the constant dark condition (DD) from day 14-28. Right panel, wavelet analysis of the wheel running activity shown in the left panel. The heatmap illustrates the strength of rhythmic activity, where warmer colors (red, yellow) indicate stronger rhythmicity. The period (in hours) is plotted on the x-axis, and the y-axis indicates successive days. **(B).** Same as in **A**, but in a *Pde10a*-KO mouse. **(C).** Average locomotor activity measured in counts per hour over a 24-hour period in control and *Pde10a*-KO mice under 12L:12D (top panel) and DD (bottom panel) conditions. Closed circles, control mice. Open circles, *Pde10a*-KO mice. Data points represent the mean ± SEM for each time point. Two-way ANOVA with Bonferroni’s *post hoc* test. (*p < 0.05, ****p < 0.0001). **D, E and F.** CREB phosphorylation analysis in control and *Pde10a*-KO mice. **(D).** Immunohistochemistry staining for phosphorylated CREB (pCREB, magenta) in the SCN at CT6 after a 15-minute light pulse in a control mouse. The SCN is marked by dashed line. Scale bars 100 µm. **(E).** Same as in **D**, but in a *Pde10a*-KO mouse. **(F).** Quantification of CREB phosphorylation in the SCN following light pulse in control and *Pde10a*-KO mice. Data are plotted as mean ± SEM (p = 0.0367, t-test). Each point represents an individual animal. **G, H and I.** pH3 phosphorylation analysis in control and *Pde10a*-KO mice. **(G).** Immunohistochemistry staining for phosphorylated H3 (pH3, cyan) in the SCN at CT6 after a 15-minute light pulse in a control mouse. The SCN is marked by dashed line. Scale bars 100 µm. **(H).** Same as in **G**, but in a *Pde10a*-KO mouse. **(I).** Quantification of H3 phosphorylation in the SCN following light pulse in control and *Pde10a*-KO mice. Data are plotted as mean ± SEM (p = 0.0167, t-test). Each point represents an individual animal. **J to O**. Behavioral phase shift in response to light in control and *Pde10a*-KO mice at CT6, CT14 and CT22. **J, L** and **M**. Actograms. Left panel, a control mouse. Right panel, a *Pde10a*-KO mouse. Animals were housed in DD. On day 8 the animal received a 1-hour light pulse (yellow bar) at the respective circadian time. The daily activity onsets of the animal are marked with the orange line, and the phase shift is visualized by the purple dashed line in each panel. **(J).** A 1-hour light pulse (yellow bar) at the CT14. **(K).** Quantification of the phase shifts observed in **J**. Data are plotted as mean ± SEM (p = 0.3571, t-test). Each point represents an individual animal. **(L).** A 1-hour light pulse (yellow bar) at the CT22. **(M).** Quantification of the phase shifts observed in **L**. Data are plotted as mean ± SEM (p = 0.0001, t-test). Each point represents an individual animal. **(N).** A 1-hour light pulse (yellow bar) at the CT6 **(O).** Quantification of the phase shifts observed in **N**. Data are plotted as mean ± SEM (p = 0.0001, t-test). Each point represents an individual animal.

### *Pde10a* inhibits CREB activation during the subjective day

We next investigated light-induced CREB phosphorylation in the SCN in the absence of *Pde10a* specifically during the subjective day, where *Pde10a* expression reaches its peak (Fig. 4C and 4D). In control mice, CREB phosphorylation was not detected after a 15-minute light pulse at CT6, consistent with previous studies(*38, 39*). Remarkably, in *Pde10a^-/-^* mice, the same light pulse at CT6 resulted in substantial CREB phosphorylation (Fig. 5D-5F). We further investigated a downstream target of CREB activation(*46*), histone-3 (H3), whose phosphorylation is associated with phase shifts of the circadian clock(*37, 40*). We observed a significant increase in H3 phosphorylation in *Pde10a-KO* mice compared to control mice following the 15-minute light pulse at CT6 (Fig. 5G-5I). These results show that in the absence of *Pde10a* expression, CREB phosphorylation is sufficient to induce downstream transcriptional and epigenetic changes in response to light stimuli during the daytime. These findings suggest that *Pde10a* gates SCN light responsiveness for circadian phase shifting by preventing CREB activation via cAMP during the subjective day.

### Light induces phase shift in *Pde10a-KO* mice during the “dead zone”

The substantial increase in CREB and H3 phosphorylation following a light pulse at CT6 in *Pde10a-KO* mice prompted us to investigate the capacity for light to induce phase shifts in these animals. We measured the phase shift in response to a light pulse in *Pde10a-*KO mice at different circadian times of the clock: CT14 (delay phase), CT22 (advance phase) and CT6 (“dead zone”). At CT14, a one-hour light pulse induced similar phase delays in both control (2.1±0.2 hours, n=6 mice) and *Pde10a-KO* mice (2.3±0.1 hours, n=4 mice; Fig. 5J-5K and S5B). At CT22, a one-hour light pulse induced phase advances in both control and *Pde10a-KO* mice, but the amplitude of the phase advance was significantly larger in *Pde10a-KO* mice (0.5±0.03 hours, n=6 mice) compared to control mice (0.2±0.04 hours, n=4 mice; Fig. 5L-5M and S5C). At CT6, the behavioral “dead zone”, a one-hour light pulse induced no phase shift in control mice (- 0.1±0.1 hours, n=8 mice), but remarkably induced a ∼1-hour phase delay in *Pde10a-*KO mice (0.8±0.2 hours, n=7 mice; Fig. 5N-5O and S5D). These results demonstrate a critical role for *Pde10a* in tuning the magnitude of phase shifts throughout the day and gating light input to the circadian clock during the daytime “dead zone”.

## DISCUSSION

In this study we reveal the underlying mechanism behind an enigma in circadian biology, namely the differential responsiveness of the suprachiasmatic nucleus (SCN) to light at different times of the day. Why does light only affect circadian behavior during the night, but not during the day? Such distinct behavioral responses to light at different circadian phases are believed to be due to the presence of a “gate” in the central circadian clock located in the SCN(*17, 47*). Transcriptomic analyses of circadian and light-induced changes of the SCN at different times of the day allowed us to identify *Pde10a* as a critical component for gating SCN responsiveness to light across the day. Consistent with the role of Pde10a in gating the light response of the clock, circadian rhythmic activity of Pde10a-KO mice is normal under constant darkness but is defective under a regular light-dark cycle. This is the first case where, despite the presence of a fully functional circadian clock and intact light input from the retina, the rhythmic expression of behavior is defective under regular light-dark conditions.

Our transcriptomic and spatial distribution analyses first allowed us to identify 10 distinct SCN neuronal subtypes across the circadian cycle that harmonize with and expand previously published reports of SCN neuronal diversity(*15, 16, 48*). SCN neurons are typically classified by their expresson of neuropeptide markers. It is worth mentioning that, among the subtypes identified in the SCN, we found a neuronal subtype expressing a non-neuropeptide marker *Mctp2*. Our analyses further allowed us to observe circadian rhythmic gene expression within each of the identified neuronal subtypes, though the extent of rhythmicity varied substantially among the subtypes. This variation suggests that different neuronal subtypes in the SCN have distinct roles in regulating various aspects of circadian rhythms. Our detailed atlas across different circadian times offers a powerful platform for future research to uncover the specific role of individual neuronal subtypes and how their rhythmic gene expression supports the functionality of the SCN.

Beyond focusing on SCN light responses at a single time point(*15, 16*), we further provide a comprehensive characterization of SCN light responses across different phases of the circadian cycle. Astonishingly, we found that light induced robust transcriptional changes at CT6, a time point that is characterized by minimal behavioral responses to light and considered to be a “dead zone” in the circadian phase response relation(*6, 14*). This finding contrasts with the relatively smaller transcriptional changes observed at CT14 and CT22, times that are associated with bonafide circadian regulation and sizeable behavioral responses to environmental light(*6, 14*). The robust transcriptional response at CT6 may suggest previously unrecognized regulatory mechanisms that are critical to prevent the SCN from shifting its clock phase during this period of the circadian cycle. A complementary hypothesis is that the SCN can effectively dissociate itself from the molecular clock in a time dependent fashion, allowing it to relay light information to the brain during the subjective day without affecting the phase of the clock. Indeed, SCN has been shown to relay light information to the brain to modulate learning and memory without altering molecular and behavioral circadian rhythms(*49, 50*).

Interestingly, although light at night elevates neuronal activity of SCN neurons(*4, 37, 51*). Our analysis revealed that light-induced transcriptional changes at CT14 and CT22 are predominantly characterized by downregulation rather than upregulation of gene expression. This observation suggests a general suppression of cellular activities on the molecular level during these circadian phases, which may be critical for the circadian clock’s ability to reset or adapt to environmental light. Furthermore, we found that most downregulated genes at CT14 and CT22 by light are related to energy and metabolism, suggesting a shift towards energy conservation in the SCN, while primarily clock-related genes are upregulated to adjust the clock phase. In contrast, many upregulated genes at CT6 in response to light are associated with neurotransmission and synapse function in SCN neurons. This further supports the idea that the SCN is competent to respond to light and relay light information during the day while the core clock remains unaffected.

A fundamental finding of this study is that cAMP phosphodiesterase *Pde10a* is a critical component for gating SCN light responsiveness across the day. *Pde10a* is well suited for this role as a “gate.” cAMP has widespread effects in cells, and previous studies have shown that cAMP can regulate the phase of the SCN clock(*10*) and that activity of the cAMP-dependent transcription factor CREB is correlated with light responsiveness of the SCN(*38, 39, 52, 53*). *Pde10a* exhibits rhythmic expression, with peak levels at CT6, intermediate at CT22, and low at CT14. In this work, we found that the phosphorylation of both CREB and histone H3 is inversely correlated with *Pde10a* expression. When *Pde10a* is high during the subjective day, PDE10A activity presumably lowers cAMP to levels that fail to trigger these phosphorylation events. Conversely, when Pde10a is low, cAMP accumulates, leading to increased phosphorylation of CREB and H3, which are both linked to phase shifts(*37, 38, 40, 53*).The increased pCREB and pH3 levels in response to light stimulation in *Pde10a-KO* mice during the daytime suggests an essential role of cAMP signaling in gating the behavioral light response, with PDE10A as a key regulator of this signaling pathway. It is worth mentioning that *Pde10a*-*KO* mice showed normal activity patterns in constant darkness, suggesting that they have an intact intrinsic circadian clock.

*Pde10a*-*KO* mice not only show light-induced phase delay at CT6, but also have unstable behavioral circadian rhythm under regular light-dark cycle. These results suggest that Pde10a is essential in tuning the light response and maintaining robust oscillation of the circadian clock under normal environmental conditions. Without *Pde10a*, the mechanisms that control light-induced phase shifts and clock synchronization are disrupted, leading to a failure to maintain a stable circadian rhythm. better biological rhythms in constant darkness than in a regular light-dark cycle.

We observed that *Pde10a* influences the magnitude of the phase shift rather than its direction, consistent with the fact that the direction of the phase shift is determined by the intrinsic properties of the molecular circadian clock network(*18, 54*). In mice where *Pde10a* is deleted, the magnitude of the phase shift is unaffected at CT14, consistent with the low levels of expression *Pde10a* across the circadian cycle at this timepoint. At CT22 when *Pde10a* levels are intermediate, light induces a more pronounced phase advance in mice lacking *Pde10a* compared to controls. These findings highlight a nuanced role for Pde10a in circadian light regulation across the day/night cycle. The enhanced phase advance at CT22 and the substantial phase delay at CT6 *Pde10a*-*KO* mice, nevertheless, lead us to speculate that in animals with shorter than 24 hours period like mice, light at the dead zone will cause phase delays, whereas, in animals with longer period like humans, advances will occur.

Another interesting feature of *Pde10a* is that it increases in response to light at all phases, although much less at CT6 when levels are already high. We reason that this could be a way to have checks on the system and to prevent an uncontrolled increase in cAMP levels that can cause major disruptions in the circadian system.

Light exposure at abnormal times, such as during shift work, when dealing with social jetlag, or while using devices late at night, has been shown to have profoundly negative effects on circadian rhythms(*2*). It is further suggested to increase the risk of various health issues including insomnia, dementia, depression, diabetes, heart attack and cancer(*2, 55, 56*). Altogether, our findings position *Pde10a* as a prime therapeutic target for circadian intervention, offering a promising route to develop strategies that address health concerns associated with circadian misalignment.

## Acknowledgements

We thank the members of the SLCR at NIMH/NIH and the Johns Hopkins Biology MouseTri-Lab for helpful discussions. We thank Bayu Sisay and Abdel Elkahloun of Microarrays and Single-Cell Genomics Core at NHGRI/NIH for performing the single nucleus sequencing.

## Funding

National Institute of Mental Health Grant MH002964 (S.H.) NIH grant EY027202 (H.Z.)

## Author Contributions

RK, MT, and SH conceptualized the study. RK. collected the sample for the sequening. MT, performed all the transcriptomic data analysis. RK, QT, and SY performed the FISH and immunohistochemistry analysis. RK, QT, SY, and HW performed behavioral analysis. SH supervised the study. RK, MT, HZ, and SH wrote the manuscript. All authors edited the manuscript.

## Competing interests

Authors declare that they have no competing interests.

## Data and Materials availability

### Lead Contact

All requests for additional information and resources should be directed to the lead contact, Dr. Samer Hattar

### Data and Code Availability

Raw and processed single nucleus RNA-sequencing data have been deposited in the Gene Expression Omnibus at GEO Accession #. GSE287917

This paper does not report original code.

Any additional data or information required to reanalyze data reported in this study are available from the corresponding author upon request.

## Supplementary Materials

Materials and Methods

Figs. S1 to S5

Data S1 to S3

## Notes

### Competing Interest Statement

The authors have declared no competing interest.

