## Supplementary information for "*Pde10a* gates light responses in the SCN to regulate circadian photoentrainment"

**The PDF file includes:**

Materials and Methods
Figs. S1 to S5
References

**Other Supplementary Materials for this manuscript include the following:**

Data S1 to S3

### Materials and Methods

#### Mice

All mice were handled in accordance with the guidelines of the Animal Care and Use Committees of the National Institute of Mental Health. Adult male mice, 3-4 months old, were used for all experiments. Animals were house under regular 12Light:12Dark cycles unless otherwise described. Wild-type mice B6129SF1/J (Jackson strain #101043) were used for the sequencing and Hiplex-FISH (or Multiplex) experiments. *Pde10a*-KO(*1, 2*) (*Pde10a*$^{-/-}$) (Jackson strain #008210) were used to study the function of *Pde10a*. Littermates that are heterozygous (*Pde10a*$^{+/-}$) or wild type (*Pde10a*$^{+/+}$) for the *Pde10a* allele were used as controls.

#### SCN dissection and nucleus isolation

Mice housed under 12L:12D cycles were kept in darkness for one day prior to tissue collection to prevent any effects from prior light exposure. A light pulse was then applied for 15 minutes either at circadian time 6 (CT6), or CT14, or CT22. 45 minutes after light exposure, mice were anesthetized with 5% isoflurane, and their brains were isolated under dim red light and placed ventral side up on an ice-cold coronal slice brain matrix (Kent Scientific). A 1 mm coronal slice was made with ice-cold razor blades positioned around the optic chiasm. This slice was laid flat on one of the blades, and hypothalamus tissue containing the SCN was collected using a 1 mm diameter sample corer (FST) and immediately placed in 500 µL of pre-chilled detergent lysis buffer in a pre-chilled Dounce homogenizer. Tissues were pooled from three mice per CT. Nuclei extractions were performed by detergent-mechanical cell lysis followed by homogenization and sucrose density gradient as described(*3*).

#### Single nucleus RNA-sequencing

Suspensions of single nuclei were loaded into a 10X Genomics Chromium controller and processed with the 3′ single-cell gene expression solution V3 (10X Genomics) according to manufacturer's instructions. All libraries were sequenced on the Illumina NextSeq 500 platform.

#### Read alignment and processing

Reads were demultiplexed and aligned with the CellRanger pipeline from 10X Genomics (v6.0.0). Read demultiplexing and alignment was performed at the Maryland Advanced Research

Computing Center, and data analysis was performed on local Mac/Linux devices using the Seurat package in R(*4, 5*).

**Integration and Clustering**

Cells with >500 unique molecular identifiers (UMIs) and <5% mitochondrial gene expression were retained for downstream analysis. For each circadian time point (CT6, CT14, CT22), light (L) and dark (D) condition, data were normalized using SCTransform(*6*) and integrated with the canonical correlation analysis (CCA) pipeline in Seurat, using the dark condition samples as the reference. This procedure resulted in three integrated datasets: CT6LD, CT14LD, and CT22LD. Principal component analysis (PCA) was performed on each dataset using 70 principal components (PCs), followed by uniform manifold approximation and projection (UMAP) for dimensionality reduction and graph-based clustering (Seurat FindClusters function). Low-quality clusters were removed based on the following criteria: (1) enrichment of >35 ribosomal protein genes ("Rpl") or >10 mitochondrial genes ("mt-"), (2) nCount_RNA or nFeature_RNA <900, and (3) a median percent.mt value >1.0. Following low-quality cluster removal, all cells across CTs were integrated into a single dataset, using the same SCTransform, CCA, 70-PC, clustering pipeline, without a reference sample. Two additional low-quality clusters were removed following this second integration based on the same filtering criteria as above.

Clusters were assigned to seven major cell types using canonical marker gene expression: astrocytes (*Agt*, *Bcan*), oligodendrocytes (*Olig1*, *Mog*), ependymal cells (*Dnah6*, *Dnah12*), microglia (*Ctss*, *C1qb*), vasculature (*Pecam1*, *Cdh5*), GABAergic neurons (*Gad1*, *Slc32a1*), and glutamatergic neurons (*Slc17a6*, *Slc17a7*). Neuronal nuclei were then subset and reprocessed using the same SCTransform, CCA, 70-PC, clustering pipeline across all conditions, without reference. Two additional clusters were excluded due to low nFeature_RNA counts, and one cluster was removed based on expression of glial markers (*Gja1*, *Agt*, *Sox6*), likely representing neuroglial doublets.

Each cluster was further assigned to a specific brain region based on manual examination of gene enrichment and specificity, using the Allen Brain Atlas ISH database and published annotations(*7*). Regions identified include the suprachiasmatic nucleus (SCN: *Vipr2, Lhx1, Vip*), medial preoptic nucleus (MPN: *Esr1, Pappa, Nts*), paraventricular nucleus (PVN: *Sim1, Ebf1,*

*Trh*), ventromedial hypothalamus (VMH: *Nr5a1, Hydin*), arcuate nucleus (ARC: *Agrp, Pomc*), lateral hypothalamic area (LHA: *Lmo7, Meis2, Foxp2*), subparaventricular zone (SBPV: *Pmfbp1, Etv1*), bed nucleus of the stria terminalis (BNST: *Moxd1*), thalamus (THAL: *Lef1*), anterior hypothalamic nucleus (AHN: *Chd7*), and anteroventral periventricular nucleus (AVPV: *C1ql2, Tnxb*).

Finally, SCN-labeled clusters were subset and processed with a final SCTransform/CCA step using 30 PCs for dimensionality reduction and clustering. Several small clusters expressing non-SCN markers and one low-quality cluster were removed, resulting in the identification of 10 distinct SCN neuronal subtypes.

**Marker gene and differential gene expression (DGE) algorithms/statistics**

The default Seurat v4 DGE algorithm was used for all single cell RNA-seq differential expression analysis (Wilcoxon rank rum, FDR-adjusted). For DGE analysis, clusters were split by 6 conditions and compared as indicated in each result/Fig..

**Gene ontology**

Gene ontology (GO) enrichment analysis was performed using the clusterProfiler R package(*8*).

**RNAscope spatial and temporal analysis**

Brain section was cut at 20-µm thickness and affixed onto microscope slides. *In situ* hybridization (Hiplex assay) were performed following a published protocol(*9*). The Hiplex-probes were from Advanced Cell Diagnostics and included *Strip2* (835581-T2), *Synpr* (500961-T4), *Calcr* (452281-T3), *Penk* (318761-T3), *Nms* (472331-T4), *Vip* (415961-T6), *Grp* (317861-T7), *Cck* (402271-T8), *Avp* (401391-T9), *Mctp2* (1072401-T1), *Bdnf* (424821-T3), *C1ql3* (495681-T5).

To visualize the *Pde10a* and *Sst* expression in the SCN, a manual Multiplex RNA-scope assay was performed following a published protocol(*10*). The probe was from Advanced Cell Diagnostics and included *Pde10a* (466201-C1), *Avp* (401391-C3), *Vip* (415961-C2), *Sst* (482691-C1).

**Immunohistochemistry (IHC)**

To examine light-induced CREB phosphorylation and histone H3 phosphorylation at CT6, mice were exposed to a 15-minute light pulse (2000 lux) and then were perfused with 4% paraformaldehyde (PFA) after 15 minutes of the light pulse. Brains were post-fixed in 4% PFA overnight and then placed in 30% sucrose in phosphate buffered saline (PBS) for two days. Brains were frozen and cut into 40 µm sections. Brain sections were first washed in 0.5% Triton X-100 in PBS (PBST). Sections were then blocked in 10% bovine serum albumin (BSA in PBST) for 1 hour and incubated overnight at 4°C in the primary antibody diluted in 2.5% BSA. Primary antibodies used were rabbit anti-pH3 mAb (1:1000; Cell Signaling Technology; Cat. No. 3377S) and rabbit anti-pCREB mAb (1:1000; Cell Signaling Technology; Cat. No. 9198S). Sections were then washed in PBST and incubated for 2 hours in the secondary antibody diluted in 2.5% BSA. Secondary antibodies used was donkey anti-rabbit Alexa Fluor 488nm (1:500; Invitrogen; Cat. No. A21206). Sections were mounted in Flouromount-G with DAPI (ThermoFisher, Cat.No. 00-4959-52).

For pH3 immunohistochemistry, the sections underwent antigen retrieval at 80 °C for 30 min in sodium citrate buffer (10 mM sodium citrate, 0.05% Tween 20, pH 6.0) before blocking in 10% BSA.

**Automated quantification for RNAscope and IHC images**

Images of brain sections were taken on a confocal microscope (Nikon, Eclipse Ti2). Images in the results are the maximal projections of a Z-stack taken through the entire 20/40 µm thick section. Any contrast or brightness adjustments were applied to the entire image in Fiji(*11*).

RNA FISH-labeled cells were counted using CellProfiler image analysis software, with an analysis pipeline modified from previous literatures(*12, 13*). Briefly, DAPI staining of nuclei was used to identify cells, and then cells with more than a set threshold number of stained speckles, or >60% of cell area covered by staining, were considered as positive for the marker. The threshold number of stained speckles varied for different markers due to various expression pattern and probe sensitivity but was kept constant for each marker across conditions.

Images of pH3 and pCREB IHC staining were also counted using CellProfiler. Because of the nucleus localization of both markers, the similar settings as used above for DAPI staining were used to identify positively stained speckles. Settings were kept consistent across conditions.

**Wheel running activity**

Animals were housed individually in cages equipped with running wheels. All mice were initially housed in a 12-hour light:12-hour dark (12L:12D) cycle, with light intensity of 2000 lux from overhead broad-spectrum LEDs, for a period of three weeks. The first week data were not used for the analysis. They were then housed in the constant dark condition (DD). Mice voluntarily ran on the wheels during their active phase at night or subjective night, with wheel revolutions recorded using Vital-View software (Mini Mitter). ClockLab software (Actimetrics) was used to visualize and calculate locomotor activity. Wavelet analysis via ClockLab was performed to assess the period under both the 12L:12D and DD conditions.

**Behavioral phase shift.**

Animals were singly housed in a cage with a running wheel under DD. The animals were exposed to bright (2000 lux) overhead broad-spectrum LED light for 1 hour either at CT6 or CT14 or CT22. Under DD, the daily onset of wheel running activity is assigned at the CT12, such that the subjective day is CT0-CT12, and the subjective night is CT12-CT24. The phase shift was calculated using ClockLab software (Actimetrics) by measuring the daily onset of wheel running activity for 7 days before and after the light pulse.

**Statistics**

The statistical tests used are listed in the results and legends. All graphs show individual animals and report the mean±SEM. Plots and analysis were done with Prism (GraphPad Prism 9.4.1).

Figure S1

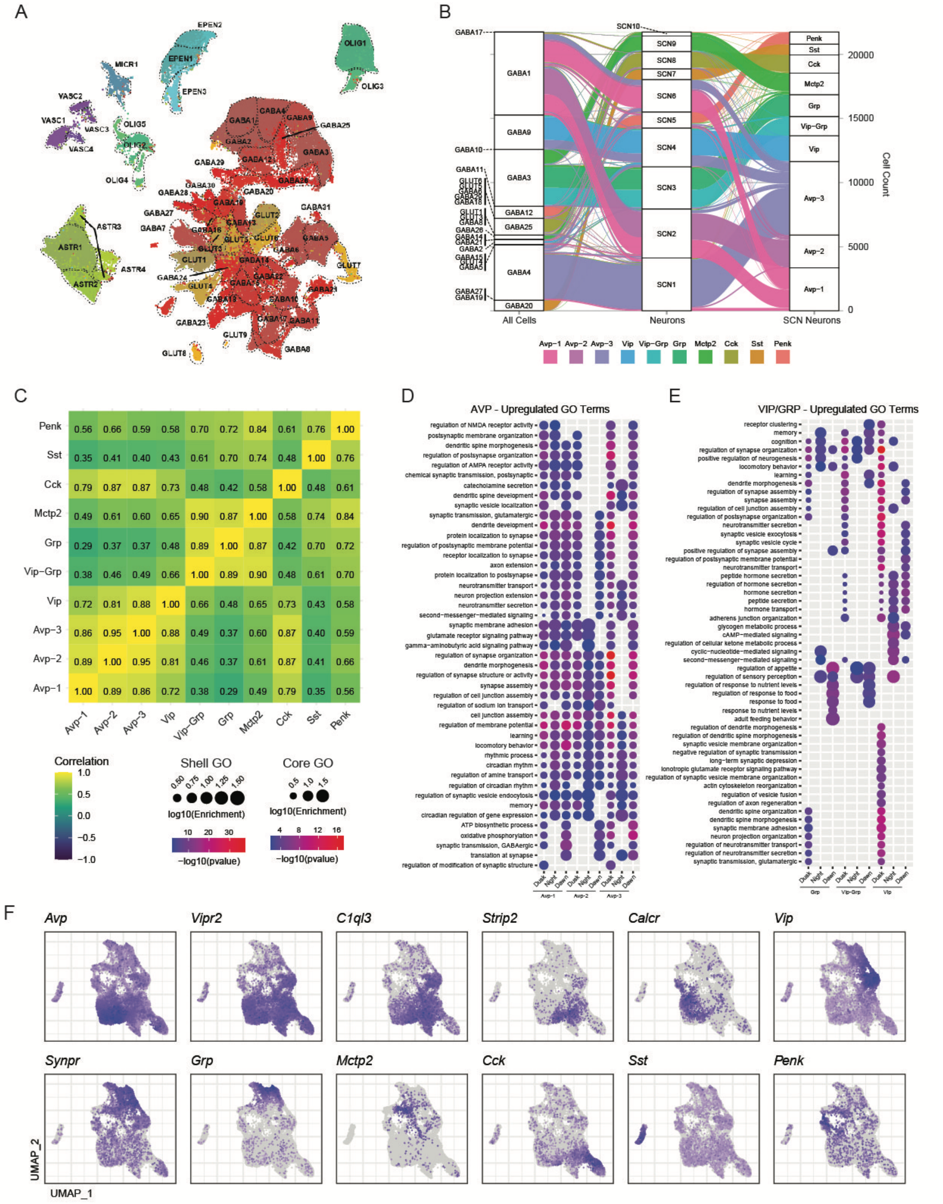


**Fig. S1. Transcriptomic profiling of SCN cells. Related to Fig. 1.**

**(A).** UMAP visualization of unsupervised clustering of 148,806 cells from the mouse anterior hypothalamus. 7 broadly defined cell types are depicted (See Materials methods). GABA: GABAergic neurons; GLUT: glutamatergic neurons; ASTR: astrocytes; OLIGO: oligodendrocytes; MICR: microglia; EPEN: ependymal cells; VASC: vascular endothelial and mural cells.

**(B).** Sankey diagram illustrating isolation and annotation of SCN neuron subpopulations. Left: major cell type annotations, as in **S1A**. Middle: intermediate hypothalamus neuron sub-class identification (see Materials Methods for details). Right: Final cluster designation, as in Fig. **1B**.
**(C).** Heatmap showing correlation coefficients (Jaccard similarity) of variable gene expression (top 2000 variable genes) between 10 distinct SCN-subtypes.
**(D).** Selected Gene Ontology (GO – biological process) term enrichment for upregulated circadian transcripts ($p < 0.05$) in Avp+ SCN neuron clusters across all CT transition comparisons: CT14D vs. CT6D (dusk), CT22D vs. CT14D (night), and CT6D vs. CT22D (dawn).
**(E).** Same as in D, but for Vip+ and Grp+ neuronal subtypes.
**(F).** UMAP visualization displaying expression of selected marker genes for 10 SCN neuronal subtypes.

Figure S2

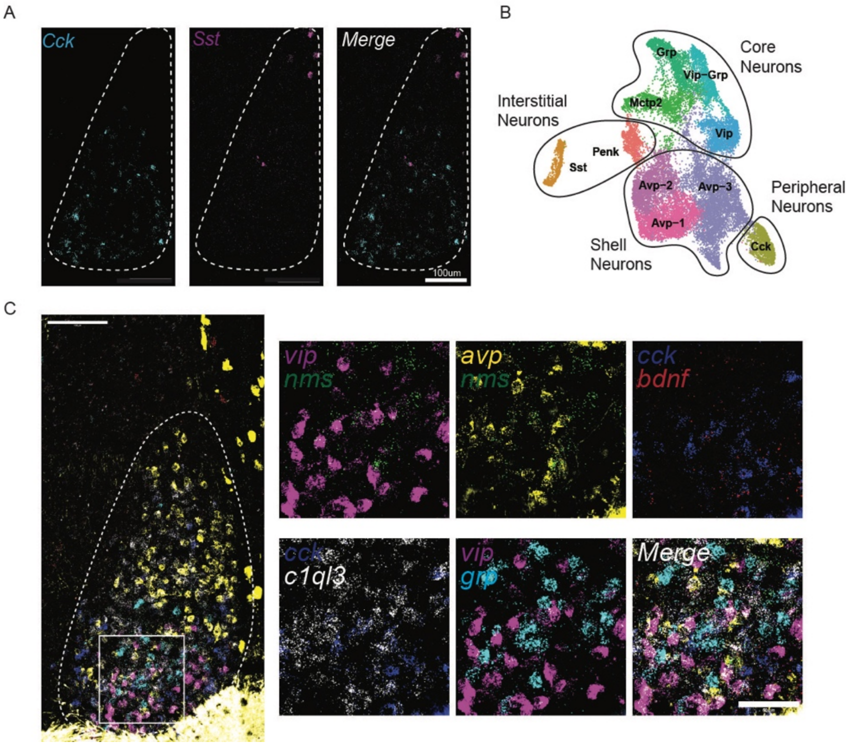


**Fig. S2. Spatial analysis of SCN neurons. Related to Fig. 2.**

**(A).** Simultaneous visualization of Cck (cyan) and Sst (magenta) neuronal subtypes in the same SCN by marker gene expression with HiPlex. Scale bar: 100 µm.

**(B).** 4 different SCN regions (core, shell, peripheral, and interstitial) depicted by the UMAP annotation of SCN neuronal subtypes.

**(C).** Left panel: Simultaneous visualization of *Avp* (yellow), *Vip* (magenta), *Grp* (cyan), *Nms* (green), *Cck* (blue), *Bdnf* (red), and *C1ql3* (white) in the same SCN by marker gene expression with HiPlex. Scale bar: 100 µm.

Right panels: magnified views of the white box marked in the left panel. From top left to bottom right: *Vip-Nms*, *Avp-Nms*, *Cck-Bdnf*, *Cck-C1ql3*, *Vip-Grp*, and merged. Scale bar: 50 µm.

Figure S3-1

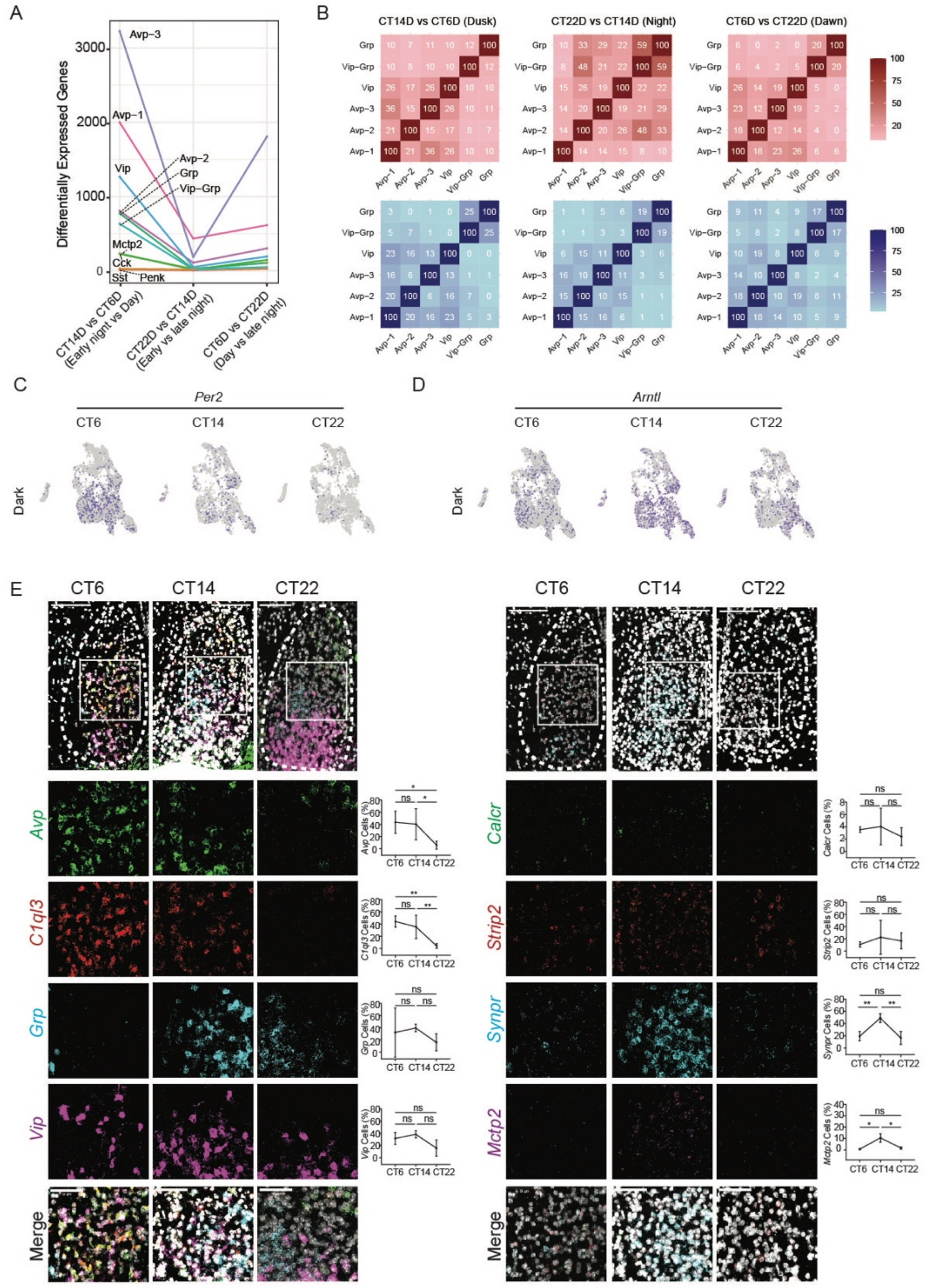


**Fig. S3-1. Temporal and spatial transcriptional changes of SCN neuronal subtypes at CT6, CT14 and CT 22. Related to Fig. 3.**

**(A).** Total counts of differentially expressed genes (DEGs) at different circadian time transitions: CT14D vs. CT6D (dusk), CT22D vs. CT14D (night), and CT6D vs. CT22D (dawn).

**(B).** Comparison matrices showing the relatedness (Jaccard similarity) of DEGs between different SCN neuronal subtypes across three distinct time periods: CT14D vs CT6D (Dusk), CT22D vs CT14D (Night), and CT6D vs CT22D (Dawn). Upper panels (red scale) and lower panels (blue scale) display relatedness between upregulated and downregulated DEGs respectively.

**(C)**. UMAP visualization displaying expression of *Per2* in dark controls at CT6, CT14, and CT22.

**(D)**. UMAP visualization displaying expression of *Arntl* in dark controls at CT6, CT14, and CT22.

**(E).** Spatial and temporal expression of SCN neuronal subtypes by HiPlex FISH at CT6, CT14 and CT22.

Left panel: top, Simultaneous visualization of *Avp* (green), *C1ql3* (red), *Grp* (cyan), *Vip* (magenta) and DAPI (white) in the same SCN with HiPlex. Scale bar: 100 µm. Lower panels, magnified views of *Avp*, *C1ql3*, *Grp*, *Vip*, and merged of the white box marked in the top panel (Scale bar: 50 µm).

Right panel: top, Simultaneous visualization of *Calcr* (green), *Strip2* (red), *Synpr* (cyan), *Mctp2* (magenta) and DAPI (white) in the same SCN with HiPlex. Scale bar: 100 µm. Lower panels, magnified views of *Calcr*, *Strip2*, *Synpr*, *Mctp2*, and merged of the white box marked in the top panel (Scale bar: 50 µm).

Graphs to the right of each marker quantify gene expression (mean ± SEM) at each time point (CT6, CT14 and CT22). Statistical significance is indicated as follows: *p* values, ns = not significant (one-way ANOVA).

Figure S3-2

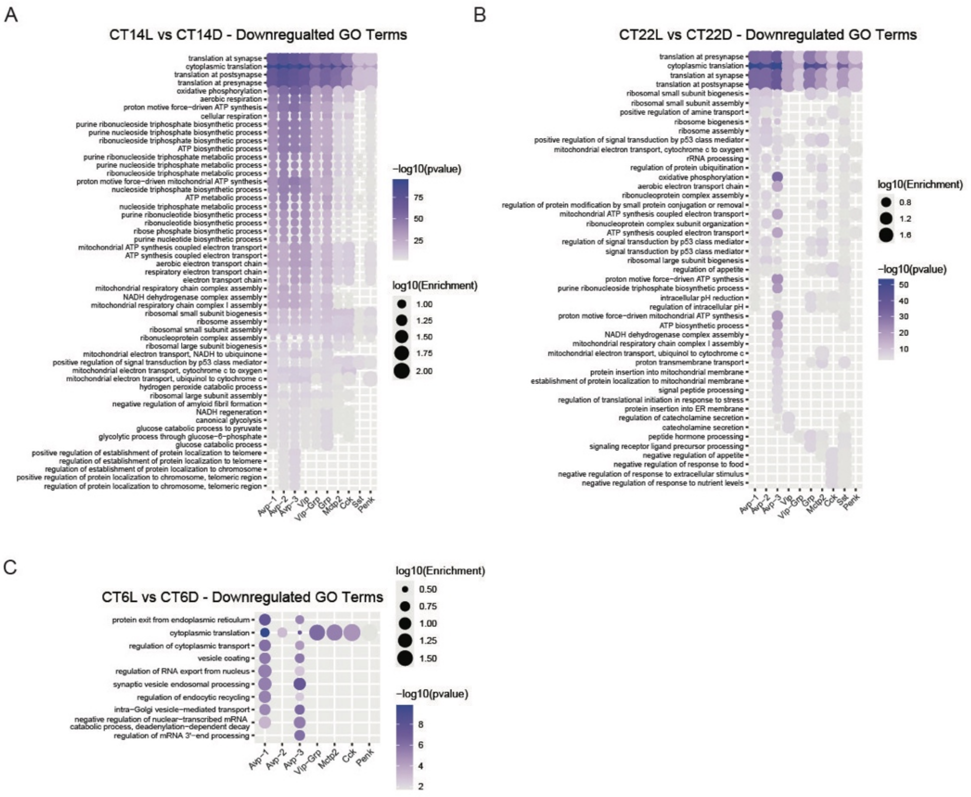


**Fig. S3-2. Light-induced transcriptional downregulation in the SCN at CT6, CT14 and CT22. Related to Fig. 3.**

**(A).** Gene Ontology (GO – biological process) term enrichment for downregulated light-induced transcripts ($p < 0.05$) in each SCN neuronal subtypes at CT14.

**(B).** Same as in A, but at CT22.

**(C).** Same as in A, but at CT6.

Figure S5

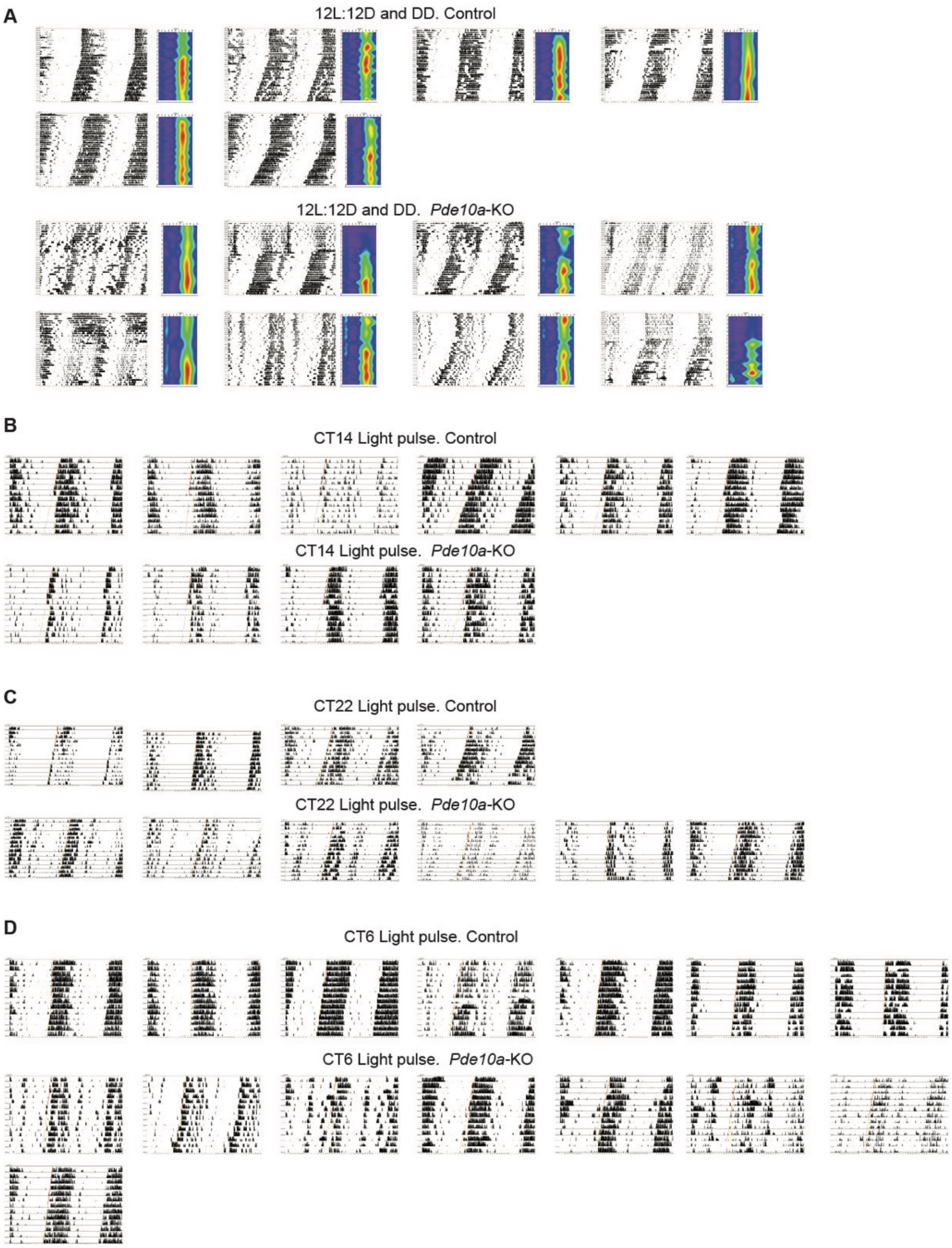


**Fig. S5. Circadian wheel running activity and behavioral phase shift in response to light in control and *Pde10a*-KO mice. Related to Fig. 5.**

**(A).** Circadian wheel running activity. The actograms of all control (upper row) and *PDE10*-KO mice (bottom row) in 12L:12D and DD conditions. Right panel of each actogram shows the wavelet analysis of the wheel running activity.

**(B).** Behavioral phase shift in response to light. The actograms of all control (upper row) and *PDE10*-KO mice (bottom row). Animals were housed in DD condition. On day 8 the animal received a 1-hour light pulse at CT14.

**(C).** Same as in B, but a 1-hour light pulse at CT22.

**(D).** Same as in B, but a 1-hour light pulse at CT6.

### Other Supplementary Materials (Separate file)

**Data S1**. Excel workbook containing quality control metrics and cell counts for all samples, related to Fig. 1.
**Data S2**. Excel workbook containing marker gene enrichment results for all SCN neuron subtypes, related to Fig. 1.
**Data S3**. Excel workbook containing differential gene expression results, related to Fig. 3 and S3-1.
